# Induction Of Chronic Stress Reveals An Interplay Of Stress Granules And TDP-43 Pathological Aggregates In Human ALS Fibroblasts And iPSC-Neurons

**DOI:** 10.1101/2020.02.17.951434

**Authors:** Antonia Ratti, Valentina Gumina, Paola Lenzi, Patrizia Bossolasco, Federica Fulceri, Clara Volpe, Donatella Bardelli, Francesca Pregnolato, AnnaMaria Maraschi, Francesco Fornai, Vincenzo Silani, Claudia Colombrita

## Abstract

Amyotrophic lateral sclerosis (ALS) and frontotemporal dementia (FTD) are fatal neurodegenerative diseases characterized by the presence of neuropathological aggregates of phosphorylated TDP-43 (P-TDP-43). The RNA-binding protein TDP-43 is also a component of stress granules (SG), cytoplasmic foci forming to arrest translation under sub-lethal stress conditions. Although commonly considered as distinct structures, a link between SG and pathological TDP-43 inclusions may occur despite evidence that TDP-43 pathology directly arises from SG is still under debate. Primary fibroblasts and iPSC-derived neurons (iPSC-N) from ALS patients carrying mutations in *TARDBP* (n=3) and *C9ORF72* (n=3) genes and from healthy controls (n=3) were exposed to oxidative stress by sodium arsenite. SG formation and cell response to stress was evaluated and quantified by immunofluorescence and electron microscopy analyses. We found that, not only an acute, but also a chronic oxidative insult, better mimicking a persistent condition of stress as in neurodegeneration, is able to induce SG formation in primary fibroblasts and iPSC-N. Importantly, only upon chronic stress, we observed TDP-43 recruitment into SG and the formation of distinct P-TDP-43 aggregates, very similar to the abnormal inclusions observed in ALS/FTD autoptic brains. Moreover, in fibroblasts, cell response to stress was different in control compared with mutant ALS cells, probably due to their different vulnerability. A quantitative analysis revealed also differences in terms of number of SG-forming cells and SG size, suggesting a different composition of foci in acute and chronic stress. In condition of prolonged stress, SG and P-TDP-43 aggregate formation was concomitant with p62 increase and autophagy dysregulation in both ALS fibroblasts and iPSC-N, as confirmed by immunofluorescence and ultrastructural analyses. We found that exposure to a chronic oxidative insult promotes the formation of both SG and P-TDP-43 aggregates in patient-derived cells, reinforcing the idea that SG fail to properly disassemble, interfering with the protein quality control system. Moreover, we obtained a disease cell model recapitulating ALS/FTD P-TDP-43 aggregates, which represents an invaluable bioassay to study TDP-43 pathology and develop therapeutic strategies aimed at disaggregating or preventing the formation of pathological inclusions.

## Introduction

Abnormal cytoplasmic aggregates of TDP-43 protein, with its concomitant nuclear clearance, represent a neuropathological hallmark of amyotrophic lateral sclerosis (ALS) and frontotemporal dementia (FTD), two fatal neurodegenerative diseases recognized as TDP-43 proteinopathies, which also share common clinical and genetic features [1–3]. Indeed, pathological inclusions of phosphorylated and ubiquitinated TDP-43 protein were found in about 97% of autoptic brain tissues from familial and sporadic ALS and in about 50% of FTD patients [3]. TDP-43 is an RNA-binding protein (RBP) that contains a C-terminal glycine-, glutamine-, asparagine-rich low-complexity domain (LCD), important for protein-protein interaction [4]. Although it localizes mainly in the nucleus, the ubiquitously expressed TDP-43 RBP is able to shuttle between the nucleus and the cytoplasm [4] where it is recruited, under stress conditions, in stress granules (SG), as demonstrated in different cellular models [5,6]. SG are reversible and dynamic membrane-less cytoplasmic protein/RNA complexes which form in response to environmental stress to temporarily arrest translation and favour only the synthesis of proteins with a pivotal cytoprotective role [7]. From a biophysical point of view, assembly of SG has been shown to occur via a liquid-liquid phase separation (LLPS) process through which soluble cytoplasmic protein/RNA complexes condensate into dynamic liquid droplets [8,9] by weak and reversible interactions of the LCDs present in several RBPs, including TDP-43 [10]. SG usually disassemble after stress removal through a chaperone-mediated mechanism, while a small fraction of aberrant SG, which accumulates defective ribosomal products (DRiPs), fails to completely disassemble and is degraded by autophagy [11–13]. Some evidence also suggest participation of the proteasome system in SG removal [9, 36], so that an interaction between chaperones, autophagy and proteasome system plays an important role in SG dynamic [14].

In the last decade, several ALS-associated proteins have been identified as components of SG in addition to TDP-43 and it has been hypothesized that SG formation participates in the neurodegenerative process, both through sequestration of RBPs and other important factors and also as an early precursor of TDP-43 pathological cytoplasmic aggregates [14,17]. In particular, when stress condition persists in the course of neurodegeneration, SG are supposed to fail to properly disassemble and may turn into insoluble and irreversible aggregates, thus disrupting neuronal homeostasis and eventually interfering with the protein quality control system [12].

Controversial reports were obtained in different studies carried out to demonstrate a co-localization of SG markers with TDP-43-positive pathological inclusions in the brain and spinal cord autoptic tissues from ALS/FTD patients [5,18–22]. Nonetheless, recent findings indicate that, even if they appear as distinct entities, a mechanistic link between SG and pathological TDP-43 inclusions may occur [23–26]. A missing point in this scenario is the evidence that TDP-43 pathology directly arises from SG, partly due to the experimental sub-lethal stress conditions used so far in several studies to induce SG formation. These short-acting and extreme insults are quite far from reproducing the persistent and subtle alterations occurring during the neurodegenerative process, which seems to be better mimicked by a mild and chronic stress condition.

In this study we provide evidence that a persistent and mild oxidative stress insult by sodium arsenite (ARS) treatment induces formation of SG in patient-derived cells, including ALS mutant fibroblasts and iPSC-derived neurons (iPSC-N), with concomitant formation of distinct cytoplasmic aggregates of phosphorylated TDP-43 (P-TDP-43). The latter hallmark is reminiscent of the pathological TDP-43 inclusions observed in ALS/FTD human autoptic brains.

## Materials and Methods

### Primary fibroblast cultures

Primary fibroblasts were isolated from skin biopsies of unrelated ALS patients carrying mutations in *TARDBP* (p.A382T, n=3) and *C9ORF72* (n=3) genes and of sex- and age-matched healthy controls (n=2), after informed consent and according to guidelines approved by the local ethics committee. Fibroblasts from 1 more healthy individual, sex- and age-matched, were obtained from Telethon Biobank. Clinical data of ALS patients, some of them already described in Onesto E. et al [27], are reported in Additional file 1: Table S1. Cells were maintained in RPMI 1640 (EuroClone, Pero, Italy) supplemented with 10% fetal bovine serum (FBS), 2mM L-glutamine, 2,5µg/ml amphotericin B, 100 units/ml penicillin and 100µg/ml streptomycin (Sigma-Aldrich, St. Louis, MO) in 5% CO_2_ incubator at 37°C. Fibroblasts were used in the experiments for no more than 12 passages.

### iPSC-derived neurons

iPSC lines from 1 healthy control, 1 mutant TDP-43 and 1 mutant C9ORF72 ALS patient were obtained by reprogramming fibroblasts (Additional file 1: Table S1) with CytoTune-iPS 2.0 Sendai Reprogramming Kit (Thermo Fisher Scientific, Waltham, MA) and differentiated into neurons, as previously described [28,29]. Briefly, iPSCs were grown in suspension in low adhesion cell culture dishes and embryoid bodies (EBs) formation was induced in 17 days by using different culture media supplemented with appropriate growth factors. EBs were then dissociated and 15,000 cells/well were plated on poly-D-Lysine/laminin (Thermo Fisher Scientific) -coated glass coverslips in 24-well plates. After additional 10 days of differentiation, the obtained iPSC-N were treated with ARS (see below). Differentiation was characterized by immunofluorescence assay with neuronal cytoskeleton markers βIII-Tubulin (1:500, Abcam, Cambridge, UK) and neurofilament (SMI-312; 1:1000, Covance, Princetown, NJ), and the MN marker HB9 (1:200, DSHB, Iowa city, IA).

### Arsenite treatment

For oxidative stress treatment, fibroblasts were exposed to 0.5mM sodium arsenite (ARS, Merck, Darmstadt, Germany) for 30 and 60 min. As no substantial differences were appreciated at 60 min in comparison with 30 min-treatment, we used the acute stress condition 0.5mM ARS for 30 min for all the following acute stress experiments. As regards chronic stress conditions, different times and different concentrations of ARS were tested, as follows: 5µM for 24-48-54-72-96 hours and 8 days; 15µM for 6-15-20-30-48 hours and 50µM for 6-16 hours. Rescue experiments were performed in fibroblasts by restoring normal growth conditions for 2 hours after acute ARS treatment and for 24-48-72-120 hours after chronic treatment. iPSC-N were exposed to 0.5mM ARS for 60 min, while for chronic stress conditions they were exposed to the following concentrations at the indicated times: 10-15µM ARS for 24-48 hours and 1-5µM for 48-72-96 hours and 7 days.

### Cell viability assay

Fibroblasts (50.000 cells/well) were plated in duplicate for each patient’s sample in 24-well dishes, 24 hours before the experiment. After treatment with 0.5mM ARS for 30 min and 15µM ARS for 30 hours, both medium and adherent fibroblasts were collected to determine viable/non-viable cells. After centrifugation at 1,000 rpm for 10 min, supernatant was discarded, and pellet was resuspended in 500µl medium. Cells were diluted 1:2 with trypan blue stain 0,4% (Logos Biosystems, South Korea) to label non-viable cells and cell viability was evaluated by using the Luna II automated cell counter (Logos Biosystems) with a custom cell-specific counting protocol.

### Immunofluorescence

Cells were fixed in 4% paraformaldehyde solution (Santa Cruz Biotechnology, Dallas, TX) for 20 min at room temperature (RT), permeabilized with cold methanol for 3 min and 0.3% Triton X-100 in phosphate buffered saline (PBS, pH 7.4, Thermo Fisher Scientific) and blocked with 10% normal goat serum (NGS, Thermo Fisher Scientific). Incubation with the primary antibodies anti-TDP-43 (1:500, Proteintech, Auckland, New Zealand), anti-TIAR (mouse 1:100, BD transduction Laboratories, Franklin Lakes, NJ; rabbit 1:300, Cell Signaling, Danvers, Massachusetts), anti-p62 (1:500, Sigma-Aldrich), anti-phosphoTDP-43 (Ser 409/410 1:200, Cosmo Bio, Carlsbad, CA) was performed in blocking solution for 1.30h at 37°C. The fluorescent-tagged secondary antibodies Alexa Fluor 488 and Alexa Fluor 555 (1:500, Thermo Fisher Scientific) were used for detection. As negative control, primary antibody was replaced by NGS. Nuclei were stained with 4’6-diamidino-2-phenylindole (DAPI) (Roche, Basilea, Switzerland) and slides were mounted with Fluorsave (Merck). Confocal images were acquired with the Eclipse Ti inverted microscope (Nikon Eclipse C1, Minato, Japan).

### Quantitative analyses of SG

Quantification of cells forming SG, of granules number and size was performed using the ImageJ software, as previously reported [30]. SG were identified by TIAR marker immunostaining and about 60 cells per sample were analyzed. On the basis of previous literature data, cells were scored as SG-positive when they presented at least two foci, three size categories of cytoplasmic granules (0.1 - 0.75µm^2^, 0.75 - 5µm^2^ and >5µm^2^) were arbitrarily considered and SG identified with a size range from 0.75 to 5µm^2^ [31–33]. Cell counting was performed in double-blind on 60x magnification images. Experiments were conducted on 3 different individuals per group of fibroblasts and for each subject the experiment was repeated 2-5 times.

### Transmission electron microscopy (TEM)

Fibroblasts (4×10^6^ cells) were incubated with the fixing solution glutaraldehyde 3% (Sigma-Aldrich) in PBS pH 7.4 for 90 min at 4°C. Cells were then collected by centrifugation at 10,000 rpm for 10 min at 4°C to form a buffy coat of cells, washed with PBS and post-fixed in 1% OsO_4_ for 1h at 4°C to prevent the formation of membranous artifacts, which may mimic autophagic vacuoles in TEM analyses [34]. Samples were dehydrated in ethanol and finally embedded in epoxy resin as previously described [35], to maintain the finest cell ultrastructure and to preserve epitopes for immunocytochemistry. Ultrathin sections (40-50 nm) were used for post-embedding immune-electron microscopy [35]. Briefly, sections were collected on nickel grids and incubated in aqueous saturated NaIO_4_ for 30 min at RT. Grids were incubated in blocking buffer (10% goat serum and 0.2% saponin in PBS) for 20 min at RT and then overnight at 4 °C in a humidified chamber with the following antibodies: LC3 (1:50, Abcam), p62 (1:20, Sigma-Aldrich), TIAR (1:20, BD transduction Laboratories) and TDP-43 (1:20, Proteintech). Grids were then washed in cold PBS before applying the gold-conjugated secondary antibodies (10 or 20 nm gold particles, BB International, Treviso, Italy), diluted (1:20) in blocking buffer for 1h at RT. Sections were examined at Jeol JEM SX 100 electron microscope (Jeol, Tokyo, Japan). Control sections were obtained by omitting the primary antibody and by incubating with only the secondary antibody.

### Quantitative analysis of TEM data

The count of 10 nm and 20 nm immune-gold particles was carried out by electron microscopy at 8.000x magnification. We scored as ATG-like vacuoles both double or multiple membranes (autophagosomes-like) containing cytoplasmic material and electron-dense membranous structures [35] A total number of 30 cells per experimental group was observed in non-serial sections of each grid. TDP-43- and P-TDP43-positive immune-gold particles were counted both in the nucleus and in the cytosol.

### Statistical analysis

Statistical analysis was conducted with PRISM software (GraphPad) by using the one-way ANOVA with Tukey’s multiple comparison post hoc test or Chi-square test for multiple groups analyses and the two-way ANOVA with Bonferroni post-tests for two-groups analyses.

## Results

### SG formation upon acute oxidative stress in mutant ALS fibroblasts

We tested whether mutations in different ALS causative genes may trigger different mechanisms of cell response to stress, since this issue has been poorly investigated so far. To study cellular response upon sub-lethal stress condition, we evaluated the formation of SG in fibroblasts from healthy controls (n=3) and ALS patients carrying mutations in *TARDBP* (p.A382T; n=3) and *C9ORF72* (n=3) genes upon high doses of ARS (0.5mM for 30 min), as previously described in literature [31]. Immunostaining performed using TIAR as a marker of SG revealed the formation of TIAR-positive cytoplasmic foci both in healthy control and ALS patient fibroblasts upon ARS treatment, but not in untreated cells, in all fibroblast groups, as expected (Fig. 1a). Interestingly, no co-localization of TDP-43 with TIAR in SG was observed in all fibroblast groups, suggesting that TDP-43 is not recruited into these ribonucleoprotein complexes following acute stress exposure in primary fibroblasts (Fig. 1a).

**Fig. 1.**
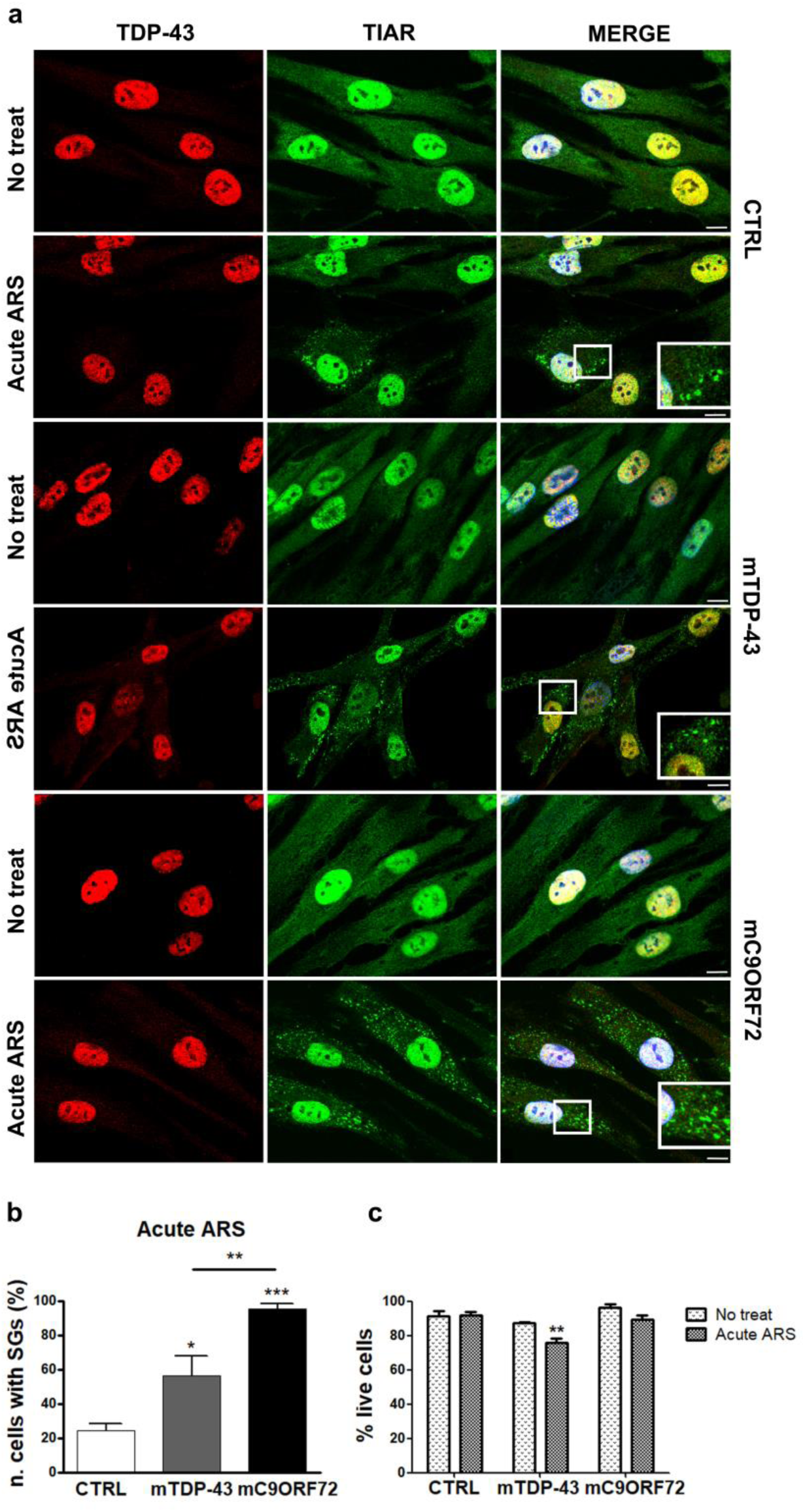
TIAR-positive SG formation upon sub-lethal stress condition in healthy control and mutant ALS fibroblasts. **a** Representative confocal images of TDP-43 (red) and the SG marker TIAR (green) in primary fibroblasts from 3 healthy controls (CTRL), 3 ALS patients carrying the p.A382T mutation in *TARDBP* gene (mTDP-43) and 3 ALS patients with pathological hexanucleotide repeat expansion in *C9ORF72* gene (mC9ORF72), before (No treat) and after acute sodium arsenite (ARS) treatment (0.5mM for 30 min). Nuclei are stained in blue (DAPI) in all merged images. Box indicates enlarged area in inset. Bar, 10µm. **b** Quantitative analysis by ImageJ software of number of cells forming SG (0.75-5µm^2^) upon acute ARS conditions in CTRL, mTDP-43 and mC9ORF72 fibroblasts (mean ± SEM, n=3 individuals per each group of fibroblasts, experiments were repeated 2-4 times for each subject and about 60 cells per sample were analyzed; one-way ANOVA with Tukey’s multiple comparison post hoc test; *p<0.05 and ***p<0.001 *vs* CTRL, **p<0.01 mC9ORF72 *vs* mTDP-43). **c** Cell viability assessed by Trypan blue stain by using LunaII instrument (mean ± SEM, n=3 individuals per each fibroblast group; two-way ANOVA with Bonferroni posttests; *p<0.05 *vs* untreated sample)

To investigate potential differences between control and mutant ALS fibroblasts in stress response and SG formation, a quantitative image analysis was performed. While only 24% control cells responded to acute ARS treatment by forming SG, mutant ALS *TARDBP* p.A382T (mTDP-43) and *C9ORF72* (mC9ORF72) fibroblasts showed higher capability to form SG, with mC9ORF72 presenting significantly more cells forming SG (95%) with respect to mTDP-43 (56%) (Fig. 1b). Inter- and intra-patient variability in SG formation is shown in Additional file 1: Figure S1. In order to evaluate toxicity induced by acute ARS treatment, we measured cell viability and found no evidence of cell death in both ARS-treated control and mC9ORF72 fibroblasts compared with the untreated samples (Fig. 1c). In contrast, a significant decreased cell viability (76%) was detected in mTDP-43 fibroblasts following acute ARS treatment in comparison with untreated cells (87%) (Fig. 1c).

As healthy control and mutant ALS fibroblasts showed a different stress response to ARS treatment in terms of number of cells forming SG, we investigated if they had also a different capability to dissolve SG upon stress removal. When we restored normal cell culture conditions for 2h after acute ARS treatment, we observed the complete disappearance of any cytoplasmic granules with no differences between control and both ALS mutant fibroblasts (Additional file 1: Figure S2).

### SG formation upon chronic oxidative stress condition in ALS fibroblasts

To investigate the hypothesis that pathological inclusions containing TDP-43 protein in ALS/FTD autoptic tissues do derive from SG, we reproduced a milder status of chronic stress *in vitro*, as it is likely to occur during the neurodegenerative process, and we evaluated if SG are able to form also in this condition and not only under short sub- lethal environmental insults as described in literature so far.

We first investigated SG formation in healthy control fibroblasts treated with low dose of ARS (15µM) in 6-48-hour time-course. We observed the formation of TIAR-positive SG at 20-hour treatment with the maximum response in terms of number of cells with SG at 30 hours (Fig. 2a). When we tested higher doses of ARS (50µM), the treatment did not induce SG formation at 6 hours, while it was toxic to cells at 16 hours.

**Fig. 2.**
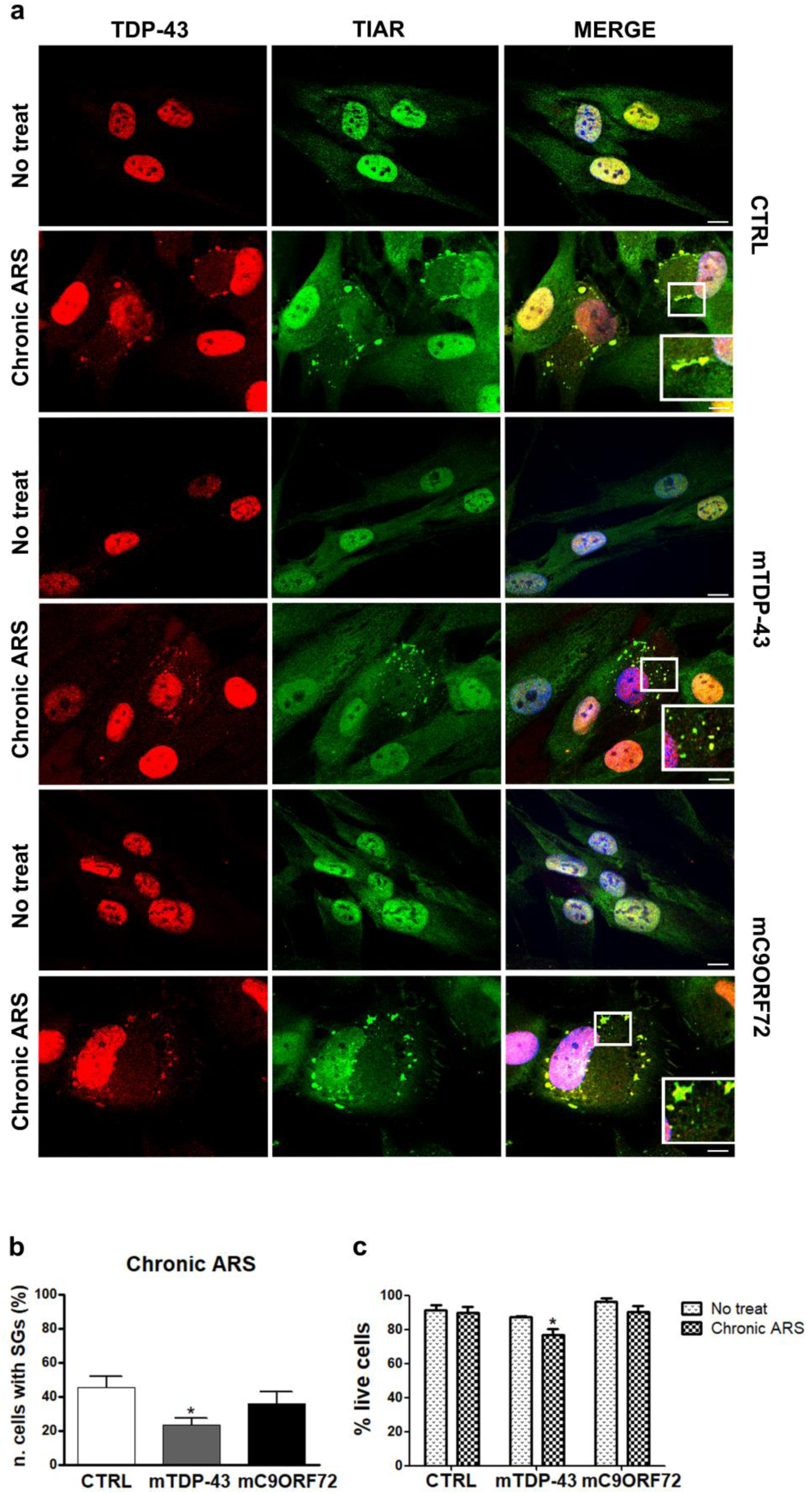
TIAR-positive SG formation upon chronic stress condition in healthy control and mTDP-43 and mC9ORF72 fibroblasts. **a** Representative confocal images of TDP-43 (red) and TIAR (green) in primary fibroblasts from CTRLs, mTDP-43 and mC9ORF72 ALS patients, before (No treat) and after chronic ARS treatment (15µM for 30h). Nuclei are stained in blue (DAPI) in all merged images. Box indicates enlarged area in inset. Bar, 10µm. **b** Quantitative analysis by ImageJ software of number of cells forming SG (0.75-5µm^2^) upon chronic ARS conditions in CTRL, mTDP-43 and mC9ORF72 fibroblasts (mean ± SEM, n=3 individuals per each fibroblast group, experiments were repeated 2-5 times for each subject and about 60 cells per sample were analyzed; one-way ANOVA with Tukey’s multiple comparison post hoc test; *p<0.05 *vs* CTRL). **c** Cell viability assessed by Trypan blue stain by using LunaII instrument (mean ± SEM, n=3 individuals per each group of fibroblasts; two-way ANOVA with Bonferroni posttests; *p<0.05 *vs* untreated sample)

In order to investigate whether mutations in ALS causative genes may influence cell response to prolonged stress in a gene-specific manner, similarly to what observed upon acute stress, we also exposed mTDP-43 and mC9ORF72 fibroblasts to chronic ARS treatment (15µM for 30h) at which we observed the maximum formation of SG in control fibroblasts. Both mutant ALS fibroblasts formed SG (Fig. 2a), although a significant lower number of SG-positive cells were observed in mTDP-43 fibroblasts (23%) compared to healthy controls (45%) (Fig. 2b). No significant differences in terms of number of cells forming SG were observed in mC9ORF72 fibroblasts (36%) compared to controls (45%), although a trend to decrease emerged from our quantitative analysis (Fig. 2b). Data showing inter- and intra-patient variability in SG formation are reported in Additional file 1: Figure S1. Interestingly, in contrast to the acute oxidative stress, we observed the recruitment of TDP-43 in SG forming in response to prolonged ARS treatment, as revealed by its co-localization with TIAR marker in all SG (100%) in both control and ALS mutated fibroblasts (Fig. 2a).

Moreover, to further prolong the oxidative stress exposure, a lower concentration of ARS (5µM) was tested on the three groups of fibroblasts for a time course ranging from 24 hours to 8 days. While 5µM ARS did not show any effect on SG formation or TDP-43 mislocalization until 96 hours, treatment for 8 days induced TDP-43 mislocalization in the cytoplasm with TDP-43-positive but TIAR-negative cytoplasmic foci, being TIAR marker largely diffused in the nucleus and the cytoplasm in both control and ALS fibroblasts (Additional file 1: Figure S3).

Similarly to acute ARS treatment, also prolonged ARS exposure did not decrease cell viability neither in healthy control nor in mC9ORF72 fibroblasts compared with the untreated samples, while treated mTDP-43 showed a significant decreased cell viability (77%) in comparison with untreated mTDP-43 fibroblasts (87%) (Fig. 2c).

Finally, we investigated whether fibroblasts were also able to disassemble SG formed in condition of chronic stress, in order to study if SG formation in condition of prolonged stress is still a reversible process. After 30-hour ARS exposure, we restored normal cell growth condition for a time-course ranging from 24 to 120 hours and, interestingly, no recovery was documented until 48 hours in all fibroblast groups. Disassembly of SG was almost complete at 72 hours and SG were completely disassembled at 120 hours with no differences between control and mutant ALS fibroblasts (Additional file 1: Figure S4).

### Differential formation of cytoplasmic granules upon acute and prolonged ARS treatment

We observed that both mutant ALS fibroblasts showed significantly less cells forming SG in response to chronic *vs* acute ARS exposure (Fig. 3a). When we quantified the amount of cytoplasmic granules, no differences were observed between control and mTDP-43 cells for the two ARS conditions (Fig. 3b). Conversely, we measured a significant increased number of granules per cell only upon acute ARS exposure in mC9ORF72 fibroblasts (32%) compared to the control group (20%), while this increase was not recapitulated after prolonged ARS treatment (18% in mC9ORF72 *vs* 19% in controls, Fig. 3b).

**Fig. 3.**
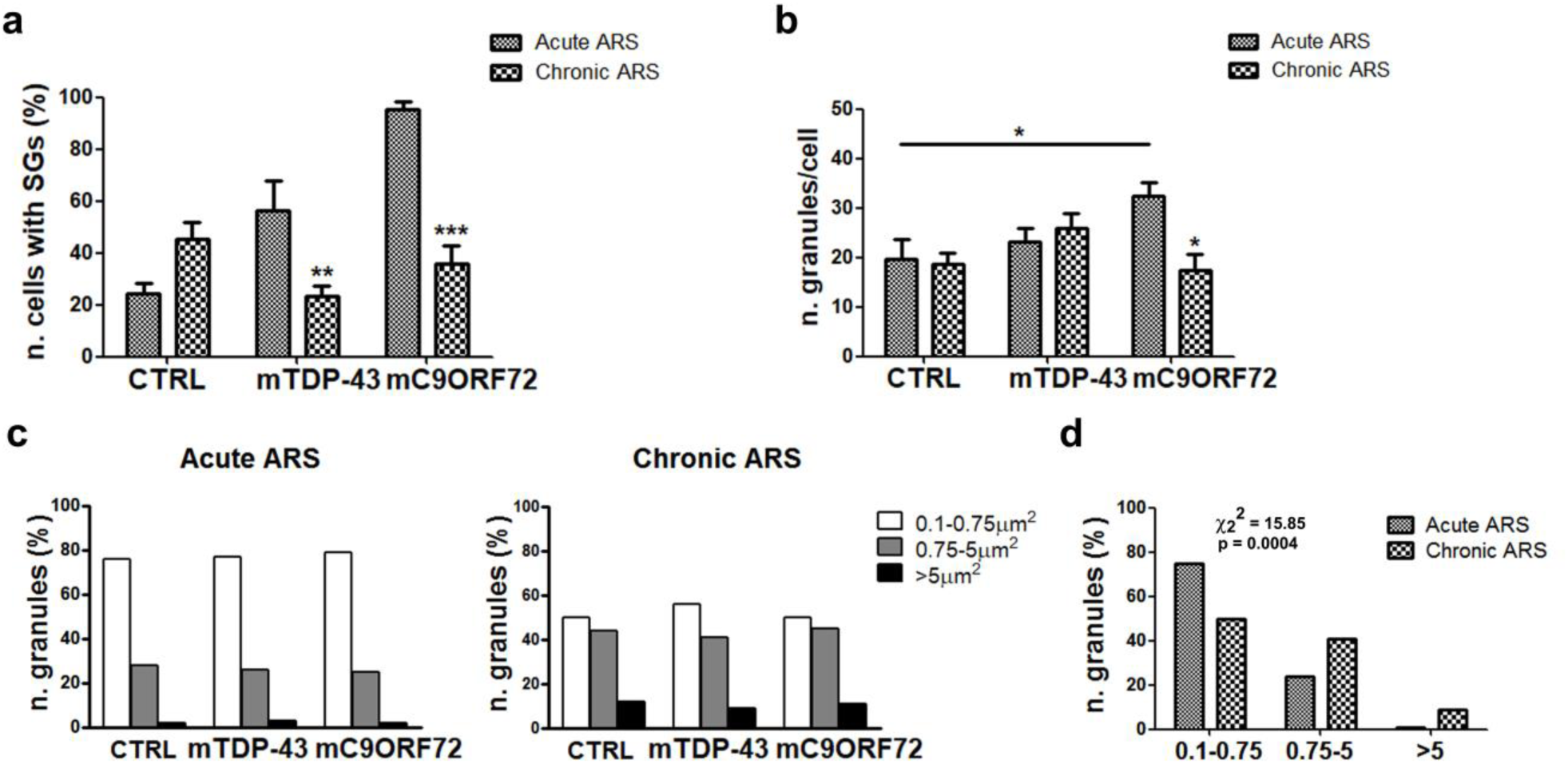
Differences in granules size and number upon acute and prolonged stress exposure. **a** Quantitative analysis by ImageJ software of number of cells forming SG (0.75-5µm^2^) upon chronic ARS conditions in comparison to acute ARS treatment in CTRL, mTDP-43 and mC9ORF72 fibroblasts (mean ± SEM, n=3 individuals per each group of fibroblasts, experiments were repeated 2-5 times for each subject and about 60 cells per sample were analyzed; two-way ANOVA with Bonferroni posttests; **p<0.01 and ***p<0.001 *vs* acute ARS treatment). **b** Quantitative image analyses by ImageJ software of granules number/cell in CTRL, mTDP-43 and mC9ORF72 fibroblasts upon acute and prolonged stress conditions (two-way ANOVA with Bonferroni posttests; *p<0.05). **c** Quantitative image analyses by ImageJ software of granules size in CTRL, mTDP-43 and mC9ORF72 fibroblasts after acute *(left panel)* and prolonged stress *(right panel)* exposure. Three interval values were arbitrarily considered (0.1 - 0.75µm^2^, 0.75 - 5µm^2^, >5µm^2^) and granules distribution in the three different classes was evaluated for each fibroblast group (Chi-square test; χ_2_^2^ = 0.5233, p = 0.97 for acute treatment; χ_2_^2^ = 1.099, p = 0.89 for chronic treatment; n=3 individuals per each fibroblast group, experiments were repeated 2-5 times for each subject and about 60 cells per sample were analyzed;). **d** Graph showing the comparison in terms of granules size distribution between acute and chronic ARS treatment. CTRL, mTDP-43 and mC9ORF72 fibroblasts were collapsed in one group as no differences emerged between the 3 groups in terms of granules size within the same stress condition (Chi square test; χ_2_^2^ = 15.85, p = 0.0004).

Our immunofluorescence analyses revealed that SG showed a bigger size after a prolonged stress condition than after acute stress exposure (Fig. 1a and 2a). We therefore performed a quantitative image analysis of cytoplasmic granules size considering three categories (0.1-0.75, 0.75-5 and >5µm^2^) where SG were identified as having a size range from 0.75 to 5µm^2^, according to previous literature data [31–33]. While no significant differences in granule size were detected among control, mTDP-43 and mC9ORF72 fibroblasts within the same stress condition (Fig. 3c), our quantitative analysis clearly showed a significant increased distribution of granules in the two categories with bigger size range (0.75-5µm^2^ and >5µm^2^) for all fibroblast groups under condition of prolonged stress (Fig. 3d), confirming an increase of SG size upon chronic oxidative stress.

### Presence of phosphorylated TDP-43 aggregates in cells exposed to chronic oxidative stress

We investigated whether TDP-43 protein recruited into SG under prolonged oxidative stress was phosphorylated at Ser 409/410, as abnormal phosphorylation and cleavage of TDP-43 C-terminal region has been reported in ALS/FTD pathological aggregates [2,36]. We found that in both physiological and acute stress conditions the P-TDP-43 protein was mainly localized in the nucleus, although mTDP-43 and mC9orf72 fibroblasts showed also a distribution in the cytoplasm of P-TDP-43-positive filaments (Fig. 4a). Moreover, after acute ARS exposure, no localization of P-TDP-43 was observed in TIAR-positive SG in both control and mutant ALS fibroblasts (Fig. 4a). However, under prolonged ARS treatment, we observed the formation of P-TDP-43-positive aggregates, with a filamentous or, more frequently, a round shape, similar to TDP-43 inclusions found in ALS/FTD autoptic brain tissues (Fig. 4a). In particular, 70% mC9ORF72 cells formed P-TDP-43 aggregates compared to 23% control and 15% mTDP-43 fibroblasts (Fig. 4b). In physiological conditions, such P-TDP-43 aggregates were present only in few cells from both control (2%) and mutant ALS patients (6% and 5% of untreated mTDP-43 and mC9ORF72 fibroblasts, respectively) (Fig. 4b). We observed a complete disassembly of these cytoplasmic aggregates after 72-hour recovery from stress with P-TDP-43 being localized in the nucleus in all fibroblast groups and not forming cytoplasmic filaments as in the original untreated ALS fibroblasts (Additional file 1: Figure S5).

**Fig. 4.**
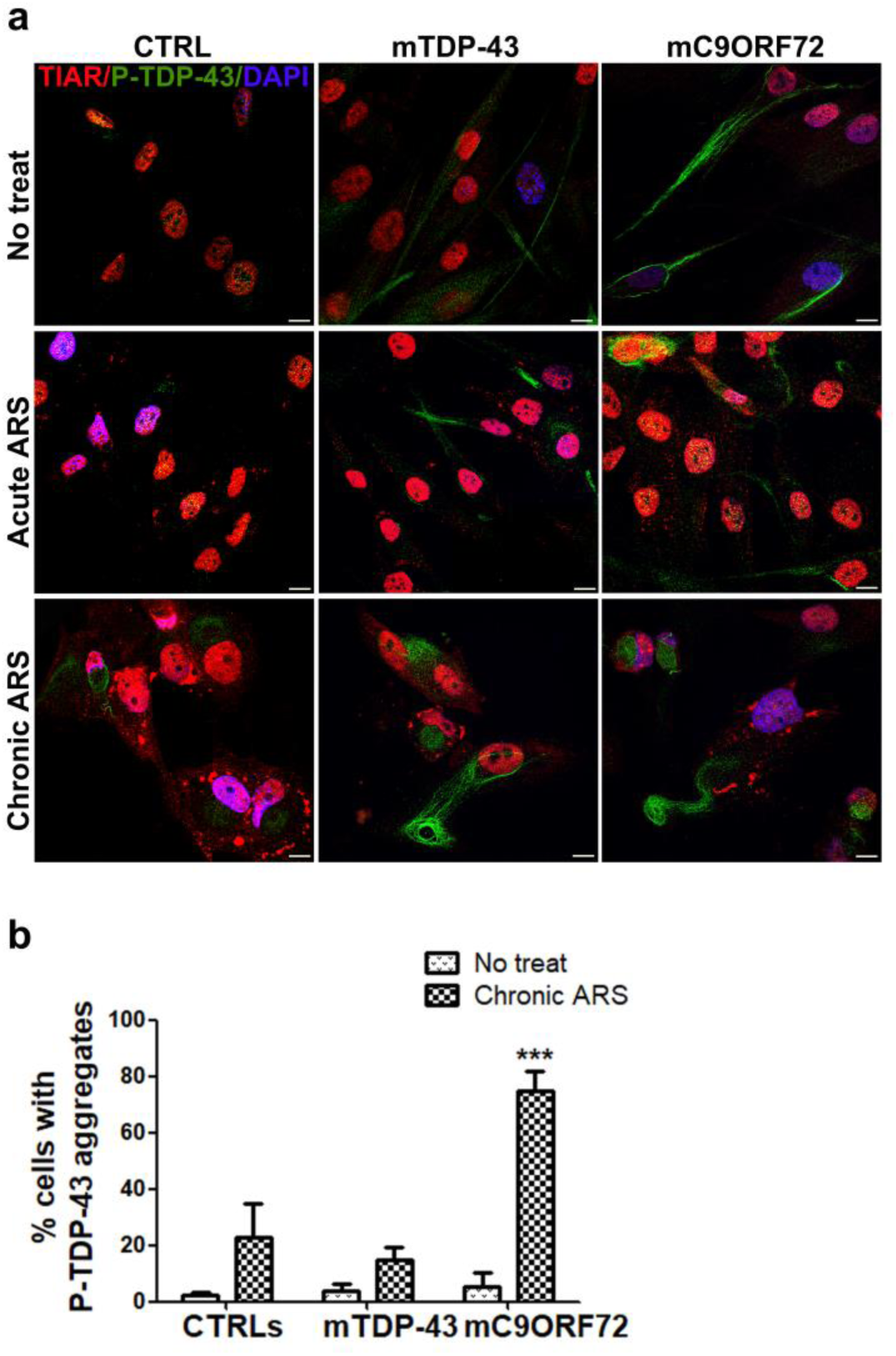
Analysis of TDP-43 phosphorylation in patients’ fibroblasts during cellular response to prolonged stress. **a** Representative confocal images of TIAR (red) and TDP-43 phosphorylated at Ser409/410 (P-TDP-43, green) in primary fibroblasts from CTRL, mTDP-43 and mC9ORF72 ALS patients, in untreated, acute (0.5mM 30min) and chronic (15µM 30h) ARS-treated cells. Nuclei are stained in blue (DAPI) in all merged images. Bar, 10µm. **b** Quantitative analysis of the number of fibroblasts (%) with P-TDP-43 aggregates in CTRL, mTDP-43 and mC9ORF72 patients (mean ± SEM, n=3 individuals per each group of fibroblasts; about 60 cells per sample were analyzed; two-way ANOVA with Bonferroni posttests; ***p<0.001 *vs* No treat and *vs* other groups).

### Impairment of the autophagy pathway upon chronic stress

The formation of TDP-43 pathological inclusions during the neurodegenerative process in ALS/FTD has been hypothesized to derive from a failure of SG to properly disassemble in condition of a prolonged insult, as a consequence of an impairment of the autophagy pathway [12]. We therefore analyzed the levels of p62, an autophagy cargo receptor used as a marker of autophagy progression. By immunofluorescence analysis, we observed low levels of p62 in both control and mutant ALS fibroblasts under physiological conditions and acute ARS treatment, while an increase of p62 in about 75% of cells was observed in all fibroblast groups upon prolonged oxidative stress (Fig. 5a). Interestingly, in this latter condition, p62 appeared mainly, but not completely, distinct from TIAR-positive SG (Fig. 5a). Following chronic stress removal, we still observed small aggregates of p62 in roughly 15% of all cells at 72-hour recovery, when P-TDP-43 aggregates completely disappeared, and persisted also at 120-hour after stress removal (Additional file 1: Figure S5).

**Fig. 5.**
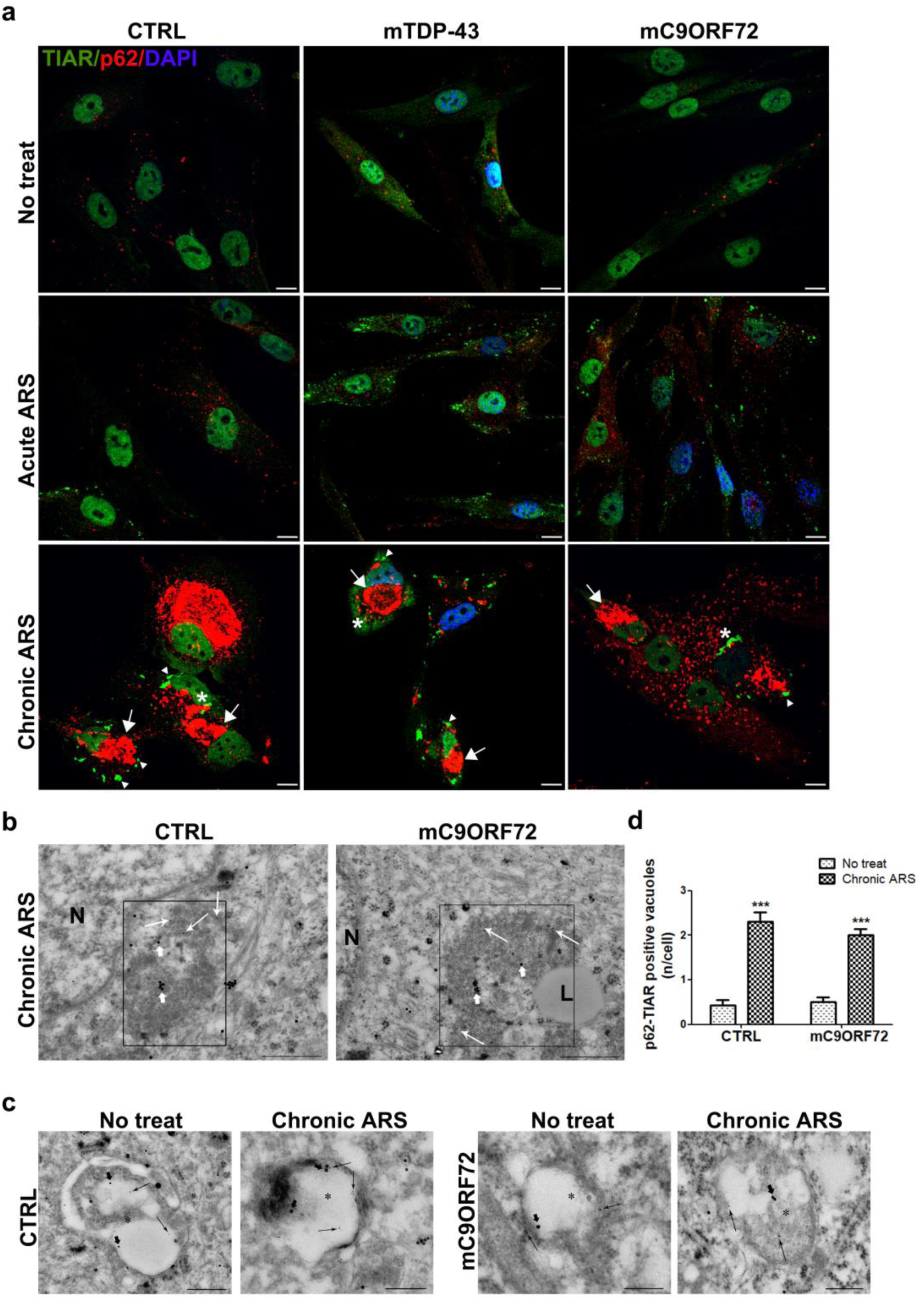
Characterization of cellular stress response by investigation of the autophagic pathway. **a** Representative confocal images of TIAR (green) and the autophagic receptor p62 (red) in primary fibroblasts from 3 CTRLs, 3 mTDP-43 and 3 mC9ORF72 ALS patients, in untreated, acute (0.5mM 30min) and chronic (15µM 30h) ARS-treated conditions. Nuclei are stained in blue (DAPI) in all merged images. Bar, 10µm. *Arrows* indicate p62 aggregates distinct from SG *(arrowheads)*; *Asterisks* indicate p62 aggregates partially co-localized with the SG marker TIAR. **b** Representative micrographs of SG in control and mC9ORF72 fibroblasts after chronic stress (15µM ARS for 30h). SG appear as cytoplasmic multi-shaped structures, not surrounded by membranes and containing fibrillar patches, stained with TIAR (10 nm Ø; *thin arrow*) and p62 immune-gold particles (20 nm Ø; *thick arrow*). Scale bar: 400 nm. **c** Representative micrographs of cytosolic autophagic vacuoles *(asterisks)* immunopositive for both p62 (20 nm Ø; *thick arrows*) and TIAR (10 nm Ø; *thin arrows*) from fibroblasts in physiological conditions and after chronic stress (15µM ARS for 30h). Scale bar: 200 nm. **d** The graph reports the number *per cell* of autophagic vacuoles positive for both p62 and TIAR in control and in mC9ORF72 fibroblasts in basal condition and after 15µM of ARS for 30h (mean ± SEM, n=30 cells for each experimental group. Two-way ANOVA; ***p<0.001 *vs* No treat)

To further investigate the connection of the autophagy pathway with SG formation in cells exposed to prolonged oxidative stress, we carried out an ultrastructural analysis as a gold standard approach to evaluate the autophagy pathway. We observed cytoplasmic multi-shaped structures, containing fibrillar patches stained with TIAR without limiting membranes and sometime also positive for p62 immune-gold particles in control and mC9ORF72 fibroblasts treated with chronic ARS (Fig. 5b), and absent in untreated cells. The involvement of autophagy pathway in SG dynamics after chronic ARS stress was evaluated by counting the number of both p62-TIAR and LC3-TIAR immune-positive autophagic vacuoles, being LC3 a marker of autophagy vesicle formation and maturation. A similar increase in the number of p62-TIAR double-stained vacuoles was observed in both ARS-treated control (2,3 vacuoles/cell) and mC9ORF72 (2 vacuoles/cell) fibroblasts compared to untreated cells (0,4 and 0,5 vacuoles/cell in control and mutant fibroblasts, respectively; Fig. 5c-d). Likewise, we detected a significantly increased number of LC3-TIAR-positive vacuoles after chronic ARS exposure in both control (1,6 *vs* 0,5 in untreated controls) and mutant fibroblasts (0,9 vs 0,5 in untreated mC9ORF72) (Additional file 1: Figure S6). The increase of double-labeled vacuoles was significantly lower in mC9ORF72 fibroblasts than in healthy control ones, supporting evidence that C9ORF72 haploinsufficiency occurring in mutant *C9ORF72* ALS patients impairs autophagy [13,37].

### Analysis of cellular response to acute and chronic oxidative stress in ALS iPSC-derived neurons

To investigate if features of cell stress response were maintained in the neuronal cells affected in the pathology, we analyzed SG formation in iPSC-neurons (iPSC-N) derived from the same ALS patients (Additional file 1: Table S1 and Additional file 1: Figure S7) in both conditions of acute and chronic oxidative stress. After acute ARS treatment we observed SG formation as already reported in literature [38] and TIAR-positive SG were present in 100% of iPSC-N (Fig. 6a), contrarily to fibroblasts (Fig. 1). In both control and mutant ALS iPSC-N, TDP-43 did not co-localize with SG (Fig. 6b), similarly to what we previously described in fibroblast cells.

**Fig. 6.**
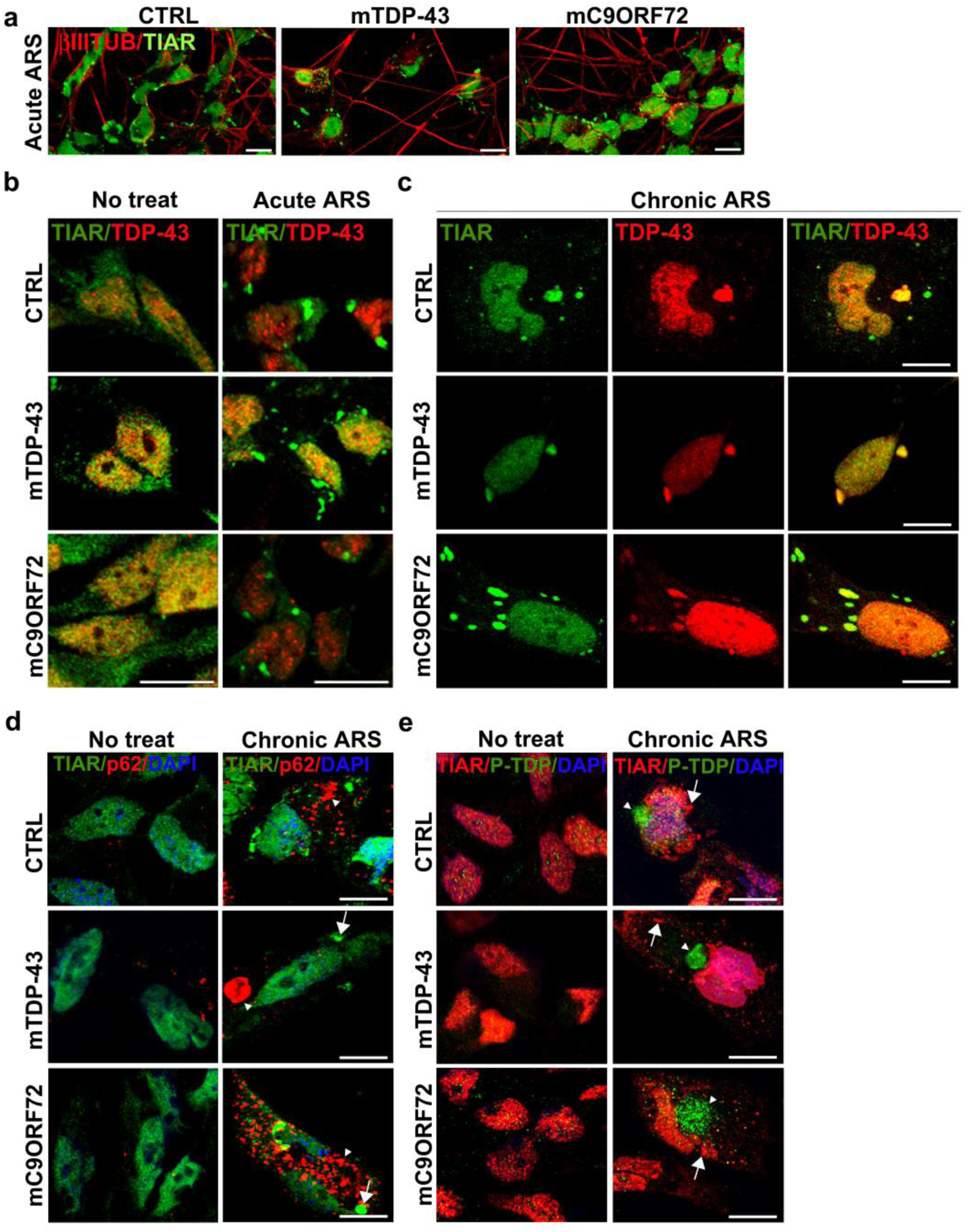
TIAR-positive SG formation and cellular response to acute and prolonged stress conditions in iPSC-N. **a** Representative confocal images of the neuronal marker βIII tubulin (red) and TIAR (green) in iPSC-N from 1 CTRL, 1 mTDP-43 and 1 mC9ORF72 ALS patient upon acute ARS treatment. Bar, 10µm. **b** Representative confocal images of TIAR (green) and TDP-43 (red) in iPSC-N from 1 CTRL, 1 mTDP-43 and 1 mC9ORF72 ALS patient before and after acute stress conditions (0.5mM ARS for 1h) and **(c)** upon chronic ARS conditions (10µM 24h). Arrow indicates SG, arrowhead indicates p62 accumulation. Bar, 10µm. Representative confocal images of **(d)** the TIAR (green) and p62 (red) and of **(e)** TIAR (red) and P-TDP-43 (Ser409/410; green) in iPSC-N from 1 CTRL, 1 mTDP-43 and 1 mC9ORF72 ALS patient before and after chronic ARS treatment (10µM 24h). Nuclei are stained in blue (DAPI) in all merged images. Arrow indicates SG, arrowhead indicates P-TDP-43 aggregates. Bar, 10µm

To set up the chronic stress condition in iPSC-N, we first exposed cells to the same chronic condition used for fibroblasts (15µM ARS for 30 hours). This treatment was toxic to iPSC-N already at 24 hour-treatment, while it was not able to induce SG formation in a shorter time interval (16 hours). Therefore, iPSC-N were subsequently exposed to lower doses of ARS (1, 5 and 10µM) and we found that a treatment with 10µM ARS for 24 hours was able to induce SG formation in 5% of cells in all three groups of iPSC-N without relevant differences between healthy controls and mutant ALS cells (Fig. 6c). Interestingly, also iPSC-N showed the recruitment of TDP-43 into SG after a prolonged stress (Fig. 6c), similarly to what we observed in primary fibroblasts (Fig. 2).

We further investigated the iPSC-N response to stress by analyzing p62 and P-TDP-43 markers upon chronic ARS treatment. Immunostaining assay showed large and abundant aggregates of p62 autophagy marker which were absent in untreated cells in both control and mutant ALS patients (Fig. 6d), confirming that, similarly to primary fibroblasts, chronic stress condition leads to a dysregulation of the autophagy pathway also in iPSC-N. As regards P-TDP-43, while in physiological condition it was mainly localized in the nucleus, under prolonged ARS treatment the phosphorylated protein formed cytoplasmic aggregates with a round shape, similar to TDP-43 inclusions previously observed in fibroblasts (Fig. 4a) and in ALS/FTD autoptic brains, in all control and mutant mTDP-43 and mC9ORF72 iPSC-N (Fig. 6e). Importantly, no co-localization of P-TDP-43 aggregates with TIAR-positive SG was observed, confirming the similar findings obtained in fibroblast cells (Fig. 4a).

## Discussion

SG have been hypothesized to be precursors of TDP-43 pathological aggregates forming during the neurodegenerative process in ALS/FTD diseases, although this issue is still under debate. In fact, almost all studies were so far carried out administering sub-lethal, short-lasting stress conditions, which markedly differ from the chronic stress conditions occurring in neurodegeneration. In addition, most studies were carried out by using immortalized cell lines showing bias when translated to patients’ cells.

In this paper, we demonstrate that a status of prolonged oxidative stress leads to formation of SG in primary fibroblasts and iPSC-N from both controls and ALS patients. Remarkably, in this chronic stress condition we also observed the recruitment of TDP-43 into SG and the presence of distinct cytoplasmic phosphorylated TDP-43 aggregates, very similar to the pathological inclusions observed in ALS/FTD brains. Moreover, an impairment of autophagy emerged from our study, suggesting that prolonged stress, as well as neurodegeneration, impairs autophagy which alters protein degradation and reduces the ability of SG to disassemble properly.

When we compared cellular response to chronic stress with to sub-lethal stress condition usually used in literature, we demonstrated that this response was different in terms of SG formation and size. In acute ARS conditions almost the totality of fibroblasts from mC9ORF72 and more than 50% of cells from mTDP-43 patients formed SG, while control fibroblasts were less responsive, suggesting that mutant ALS fibroblasts are more vulnerable to stress due to an increased susceptibility. Literature data support our results, as in condition of acute oxidative stress ALS-linked mutations were previously described to enhance TDP-43 and FUS cytoplasmic translocation and SG formation in both human cell lines and animal models [18,23,39–42]. On the contrary, a previous study on mTDP-43 fibroblasts showed that the same TDP-43 p.A382T mutation caused a significant decrease of number of cells forming SG in condition of acute stress [31]. Our study included a bigger sample size and, importantly, we showed also that inter- and intra-patient variability was particularly relevant in response to acute stress in mTDP-43 fibroblasts. We therefore can not exclude that both variability and different clinical data of patients may influence the different results obtained and we think that functional studies in patient-derived cells need more cases to be investigated. In contrast to acute stress conditions, when we reproduced a milder and prolonged status of chronic stress *in vitro*, fibroblasts from mTDP-43 patients formed less SG in comparison with control fibroblasts. We speculate that, when the stress persists becoming chronic as in neurodegeneration, mTDP-43 cells, in contrast to mC9ORF72, show less ability to induce a long-term protective mechanism. According to this hypothesis, also cell viability is specifically reduced in mTDP-43 fibroblasts upon both acute and chronic stress exposure.

Importantly, we showed for the first time that, regardless of the ALS mutation carried, TDP-43 is recruited into SG only in response to chronic, but not to acute, ARS condition in patient-derived peripheral and iPS-differentiated neuronal cells. This is in contrast to what we and other groups previously described in immortalized cell lines where localization of TDP-43 in SG occurred under sub-lethal stress conditions [5, 11]. The different response to stress of primary and immortalized cells is also supported by a recent paper showing that TDP-43 is recruited into SG after 24-hour puromycin treatment in mTDP-43 N352S and FUS R521G iPSC-motoneurons [26].

A significant increase of SG size emerged in both control and mutant ALS fibroblasts in chronic compared with acute treatment, suggesting that, after a prolonged stress, both a fusion of smaller SG and a different composition of SG may occur, as already hypothesized [14], while no differences among the fibroblast groups were observed within the acute ARS conditions, as also confirmed by Orru’ et al [31], or within the chronic ARS exposure. An altered composition of SG and an increase of cytoplasmic TDP-43 protein, as described for ALS-linked mutations, may promote interactions between the LCD of several RBPs, aberrant phase transition and aggregation, finally triggering ALS/FTD pathogenesis. In this scenario a crosstalk between SG components and aggregates may occur and several SG components may be trapped in pathological TDP-43 inclusions. This interplay may be possible because of the biophysical properties of SG, as their assembly occurs via the liquid-liquid phase separation (LLPS) process from cytoplasm [8,9] and is mediated by the LCD contained in most RBPs, such as TDP-43 and FUS, but also TIA-1, hnRNPA1, hnRNPA2, EWSR1 and TAF15 [10,43]. It has been recently shown that self-oligomerization of the LCD, an essential domain for TDP-43 recruitment into SG [5], mediates the phase transition of TDP-43 RBP in liquid droplets [23]. Despite their physiological role in the LLPS, the LCD of RBPs may also favour a liquid-to-solid transition and, consequently, be responsible for protein misfolding and aberrant protein aggregation of RBPs observed in several neurodegenerative diseases [9,25,40,44–46].

By reproducing a status of chronic oxidative stress with low concentration of ARS for a prolonged time in ALS patient-derived cells, we observed a parallel formation of TIAR/TDP-43-positive SG and of distinct phosphorylated TDP-43-positive aggregates, not localizing with SG. In support of our data, two recent studies, by using two different approaches in mammalian cell lines, showed formation of granules of phosphorylated and ubiquitinated TDP-43, distinct from SG, suggesting that TDP-43 inclusions may form through a SG-independent pathway [23,24]. Moreover, during acute ARS exposure these studies demonstrated that cytoplasmic TDP-43 underwent transition from liquid droplets into gel/solid-like structures, while the TDP-43 protein inside SG remained in a dynamic state within liquid-like RNA-rich droplets. Therefore, these studies do not exclude the possibility that pathological TDP-43 granules may also evolve through an intermediate transition from SG and our results also support this possibility. According to our results, a recent study also demonstrated that TDP-43 protein recruited into SG in immortalized cell lines was not phosphorylated and, after treating cells with ARS 25µM up to 6h, formation of P-TDP-43-positive foci was observed, although SG were absent [25]. Altogether, these results may help explain why several studies were not able to identify SG markers in pathological TDP-43 inclusions in post-mortem ALS/FTD brain tissues [5,19,21].

Consistent with our findings, while TDP-43 is protected by phosphorylation when it is recruited within SG, as previously hypothesized [23,25], when the stress persists becoming chronic as in neurodegeneration, cytoplasmic full-length TDP-43 protein may be in part released from SG and undergo phosphorylation by protein kinases, such as GADD34/CK1ε [47]. This may trigger formation of P-TDP-43 inclusions, driving a pathological aggregate transition and leading to an engulfment of the autophagy and other systems involved in protein quality control. It was shown that, when autophagy is impaired, SG disassembly is also affected because of accumulation of misfolded proteins including DRiPs [11], producing insoluble aggregates that disrupt neuronal homeostasis and promote ALS pathogenesis and neurodegeneration [12]. Indeed, defects in SG assembly and abnormal inclusions of TDP-43 protein have been described not only in ALS and FTD, but also in other neurodegenerative diseases, including spinocerebellar ataxia, Alzheimer’s, Parkinson’s and Huntington diseases [48], suggesting that TDP-43 may have a common pivotal role in neurodegeneration.

Our data also show that P-TDP-43 aggregates dissolve with SG during recovery from a prolonged stress so that we speculate that in neurodegeneration a critical stress threshold exists over which SG disassembly becomes an irreversible process and causes an engulfment of the protein quality control system, including chaperones, and the autophagic and ubiquitin/proteasome systems. Interestingly, we observed p62-positive aggregates after prolonged ARS treatment, suggesting autophagy impairment [49]. Additionally, our electron microscopy analysis shows that chronic ARS treatment increases the number of p62-TIAR- and LC3-TIAR-positive autophagic vacuoles, reinforcing the idea that p62, through autophagy, plays a direct and pivotal role in SG clearance. While P-TDP-43-positive aggregates completely dissolved at 72-hour recovery from prolonged ARS treatment, smaller p62 aggregates persisted at 120 hours, confirming that the autophagy system may be impaired as an initial and persistent event.

Interestingly, upon prolonged oxidative stress almost all the mutant C9ORF72 fibroblasts formed filamentous and round shapes P-TDP-43 inclusions in comparison with control and mTDP-43 fibroblasts, supporting an important involvement of C9ORF72, a regulator of endosomal and vesicular trafficking, in autophagy [50]. Remarkably, it has been demonstrated that C9ORF72 colocalizes with Ubiquilin-2 and LC3-positive vesicles and the ratio of the autophagosome markers LC3II/LC3I is altered in *C9ORF72* knocked-down cells [51]. Additionally, C9ORF72 protein was recently described to form a complex with the autophagy receptor p62, which associates with symmetrically methylated arginine proteins enriched in SG, controlling SG removal by autophagy [13]. All these data, together with present findings, suggest that an impairment of the p62-dependent clearance of SG components may be amplified in ALS patients with *C9ORF72* repeat expansions, due to the haploinsufficiency of C9ORF72 protein observed in such patients [52].

## Conclusions

We proved, for the first time, that formation of SG is induced also by a mild and chronic oxidative stress in primary fibroblasts and iPSC-N with differences in stress response between healthy controls and ALS patients, in association to the appearance of distinct P-TDP-43 cytoplasmic inclusions. As the transition of ALS-linked RBPs from a liquid-phase separation to solid aggregates was shown to be time-dependent [53], we here hypothesize that a prolonged ARS treatment is more likely to induce irreversible aggregation. Therefore, ALS patient-derived cells, exposed to a persistent oxidative stress, represent a suitable bioassay to study not only TDP-43 pathology, but also to test potential drugs able to prevent or disaggregate P-TDP-43 inclusions. Since most of the cellular and animal models used so far failed to fully reproduce ALS/FTD pathology and phenotype, our findings strongly sustain the translatability of pathomechanisms from patient-derived peripheral cell models such as fibroblasts to human iPSC-N, opening the scenario of new therapeutic approaches also for individual rehabilitation of patients.

## Abbreviations

ALS: amyotrophic lateral sclerosis
ARS: sodium arsenite
DRiPs: defective ribosomal products
FTD: frontotemporal dementia
iPSC-N: iPSC-derived neurons
LCD: low-complexity domain
LLPS: liquid-liquid phase separation
mC9ORF72: mutant C9ORF72
mTDP-43: mutant TARDBP p.A382T
P-TDP-43: phosphorylated
TDP-43; RBP: RNA-binding protein
SG: stress granules
TEM: Transmission electron microscopy.

## Declarations

### Ethics approval and consent to participate

Skin biopsies of healthy individuals and patients were obtained after written informed consent and in accordance with guidelines approved by the local ethics committee (IRCCS Istituto Auxologico Italiano Review Board).

### Consent for publication

Not applicable.

### Availability of data and material

Not applicable.

### Authors’ contributions

A.R. performed experiments, contributed to experimental design and data analyses and wrote the manuscript; V.G performed part of the experiments and critically revised the manuscript; P.L. performed electron microscopy experiments; P.B. generated and characterized iPSC cell lines; F.F. performed electron microscopy experiments; C.V. differentiated iPSC-N; D.B. differentiated iPSC-N; F.P. contributed to statistical analyses; A.M. performed part of the experiments on fibroblasts; F.F. critically discussed electron microscopy data and revised the manuscript; V.S. critically discussed all data and revised the manuscript; C.C. designed the experiments, elaborated results and wrote the manuscript. All authors read and approved the final manuscript.

## Acknowledgments

Special thanks to the patients because without their generous contribution this study would not have been possible. We also thank the Cell line and DNA bank of pediatric movement disorders and mitochondrial diseases of the Telethon Network of Genetic Biobanks (Project GTB07001) for donation of one healthy fibroblast cell line.

