## Supplementary data for "Induction Of Chronic Stress Reveals An Interplay Of Stress Granules And TDP-43 Pathological Aggregates In Human ALS Fibroblasts And iPSC-Neurons"

**Supplementary Table S1** Clinical features of healthy controls, *TARDBP*- and *C9ORF72*-mutated patients.

| Patient ID | Sex | Age at biopsy | Gene | Mutation | Diagnosis | Family history | Age at onset | Site of onset | Cognitive impairment | Available iPSCs |
|---|---|---|---|---|---|---|---|---|---|---|
| **C1** | M | 37 | - | - | - | - | - | - | - | - |
| **C2** | F | 45 | - | - | - | - | - | - | - | Yes |
| **C3** | F | 53 | - | - | - | - | - | - | - | - |
| **T1** | M | 26 | TARDBP | p.A382T | ALS | No | 25 | spinal | No | - |
| **T2** | M | 56 | TARDBP | p.A382T | ALS | No | 54 | spinal | No | Yes |
| **T3** | F | 60 | TARDBP | p.A382T | ALS | No | 59 | spinal | No | - |
| **C9-1** | M | 45 | C9ORF72 | 1500 units | ALS | No | 44 | spinal | No | - |
| **C9-2** | F | 48 | C9ORF72 | 2200 units | ALS | No | 47 | spinal | No | Yes |
| **C9-3** | M | 64 | C9ORF72 | 1200 units | ALS | No | 61 | spinal | No | - |

**Supplementary Fig. S1** TIAR-positive stress granules (SG) formation upon sub-lethal and prolonged stress condition in healthy control and mutant TDP-43 and C9ORF72 fibroblasts. Quantitative analysis by ImageJ software of number of cells forming SGs (0.75-5µm$^2$) upon acute and chronic ARS conditions in CTRL, mTDP-43 and mC9ORF72 fibroblasts. Inter- and intra-patient variability is shown in the graph (mean ± SEM, n=3 individuals per each fibroblast group; acute and chronic stress experiments were repeated 2-4 times and 2-5 times respectively for each subject. About 60 cells per sample were analyzed; one-way ANOVA with Tukey's multiple comparison post hoc test; *p<0.05 and ***p<0.001 *vs* CTRL, **p<0.01 mC9ORF72 *vs* mTDP-43).

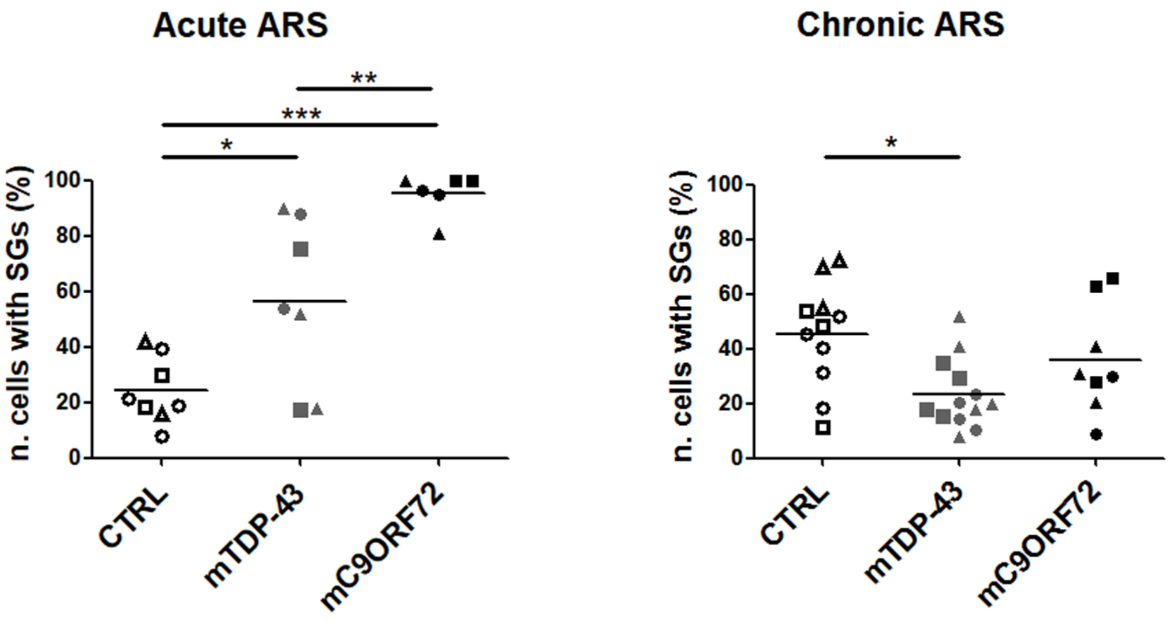

**Supplementary Fig. S2** Recovery experiments from acute ARS treatment. Representative confocal images of TDP-43 (red) and the SG marker TIAR (green) in primary fibroblasts from 3 CTRL, 3 mTDP-43 (p.A382T) and 3 mC9ORF72 ALS patients, in condition of acute ARS treatment (0.5mM for 30 min) and after 2h rescue. When ARS was removed, and normal cell culture conditions restored by replacing medium containing ARS reagent with the normal growth medium for 2h, SGs disappeared in both control and mutant fibroblasts. Nuclei are stained in blue (DAPI) in all merged images. Bar, 10µm.

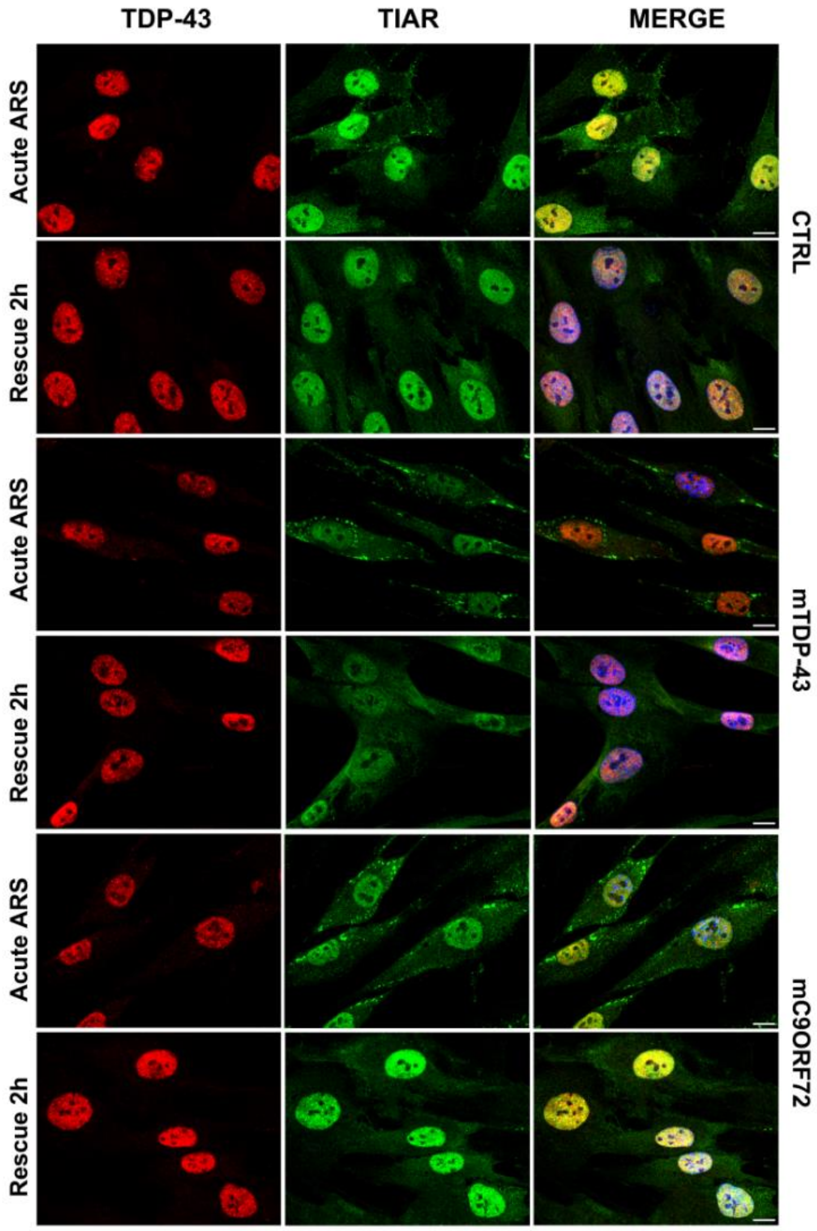

**Supplementary Fig. S3** Stress response of fibroblasts to prolonged ARS treatment for 8 days. Representative confocal images of TDP-43 (red) and the SG marker TIAR (green) in primary fibroblasts from 3 CTRL, 3 mTDP-43 and 3 mC9ORF72 ALS patients, before (No treat) and after prolonged ARS treatment (5μM for 8 days). Nuclei are stained in blue (DAPI) in all merged images. *Arrows* indicate TDP-43 puncta which are negative for TIAR. Bar, 10μm.

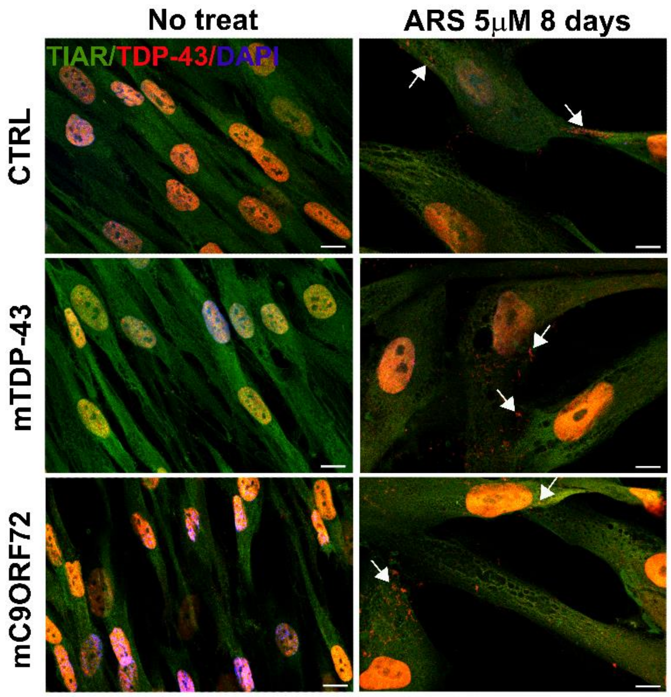

**Supplementary Fig. S4** Recovery experiments from chronic ARS treatment. Representative confocal images of TDP-43 (red) and the SG marker TIAR (green) in primary fibroblasts from 3 healthy CTRL, 3 mTDP-43 and 3 mC9ORF72 ALS patients, in condition of chronic ARS treatment (15µM for 30h) and in condition of recovery for 120h. When ARS was removed, and normal cell culture conditions restored by replacing medium containing ARS reagent with the normal growth medium for 120h, SGs disappeared in both control and mutant fibroblasts. Nuclei are stained in blue (DAPI) in all merged images. Bar, 10µm.

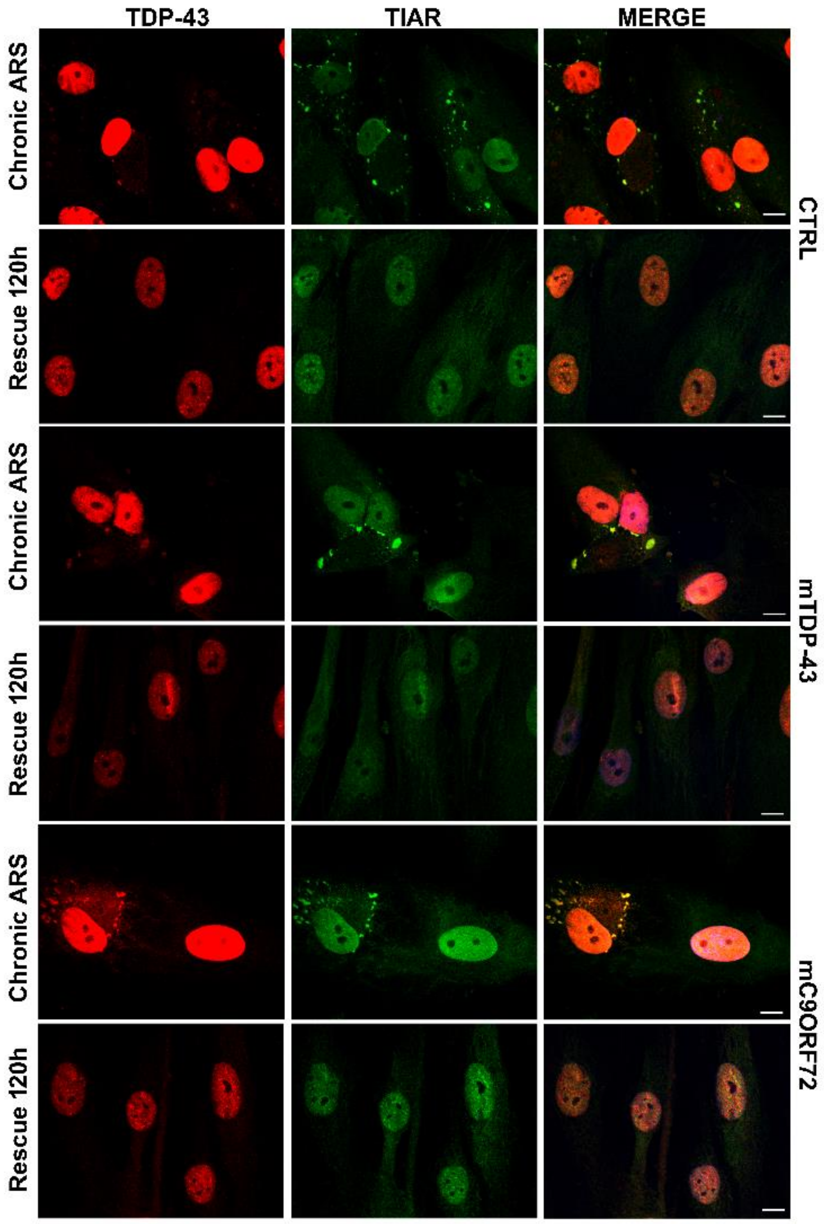

**Supplementary Fig. S5** Rescue experiments in controls and mutant ALS fibroblasts. Representative confocal images of SG marker TIAR (red) and P-TDP-43 (green) *(upper panels)* and of TIAR (green) and p62 (red) *(lower panels)* in primary fibroblasts from 3 CTRL, 3 mTDP-43 and 3 mC9ORF72 ALS patients in condition of 72-hour recovery from chronic ARS exposure. Nuclei are stained in blue (DAPI) in all merged images. Bar, 10μm.

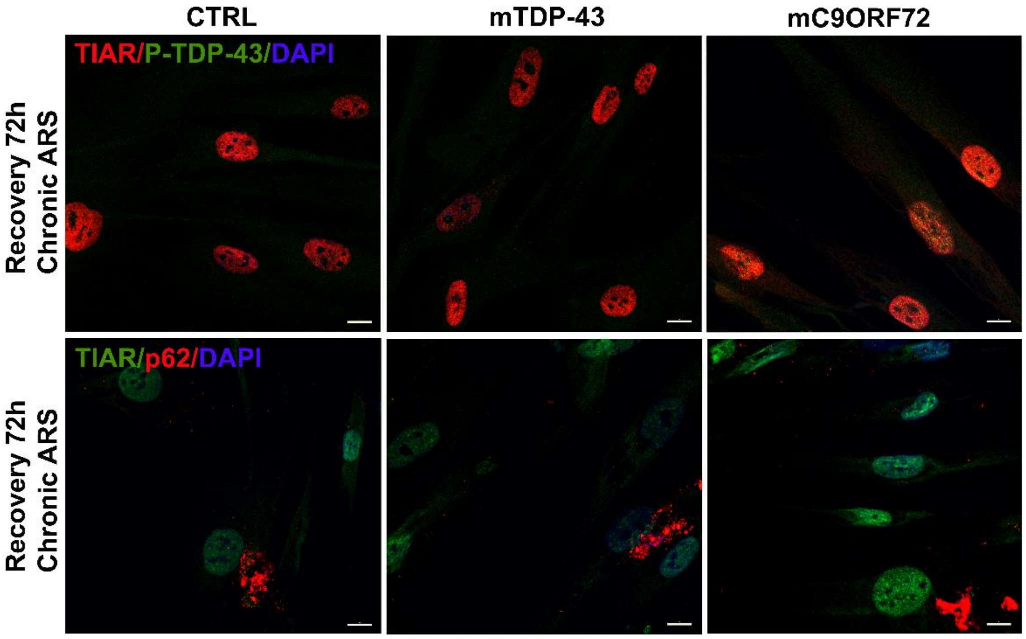

**Supplementary Fig. S6** Ultrastructural analyses of autophagy response after chronic ARS treatment in control and in ALS mC9ORF72 fibroblasts. Representative micrographs of cytosolic vacuoles *(asterisks)* immunopositive for both LC3 (20 nm Ø; *thick arrow*) and TIAR (10 nm Ø; *thin arrow*) from control and mC9ORF72 fibroblasts in physiological conditions and after chronic stress (15 µM ARS for 30h) Scale bar: 200 nm. The graph reports the number *per cell* of vacuoles immunopositive for both LC3 and TIAR. (mean ± SEM, n=30 cells for each experimental group; **p<0.01 mC9ORF72 *vs* CTRL; ***p<0.001 *vs* No treat).

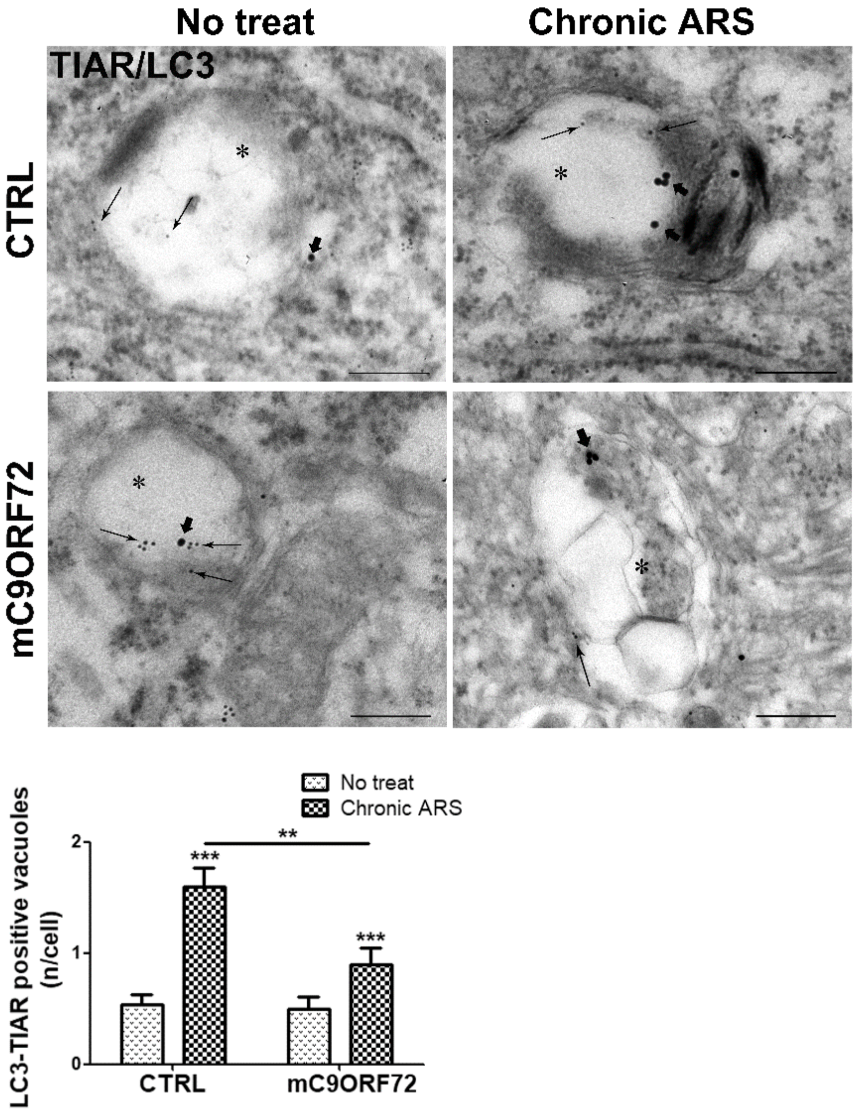

**Supplementary Fig. S7** Characterization of iPSC-N differentiated from control and mutant ALS iPSC. Representative confocal images of **(a)** βIII tubulin (green) and SMI312 (red) neuronal markers and of **(b)** βIII tubulin (green) and HB9 (red) motoneuronal marker in iPSC-N from 1 healthy CTRL, 1 mTDP-43 and 1 mC9ORF72 ALS patient. Nuclei are stained in blue (DAPI) in merged images in **(b)**. Bar, 10µm. Both control and mutant ALS iPSC-N showed neuronal processes with nearly 100% and 80% of cells positive for βIII-tubulin and SMI312 markers, respectively, while about 20% of cells stained positive for HB9 marker.

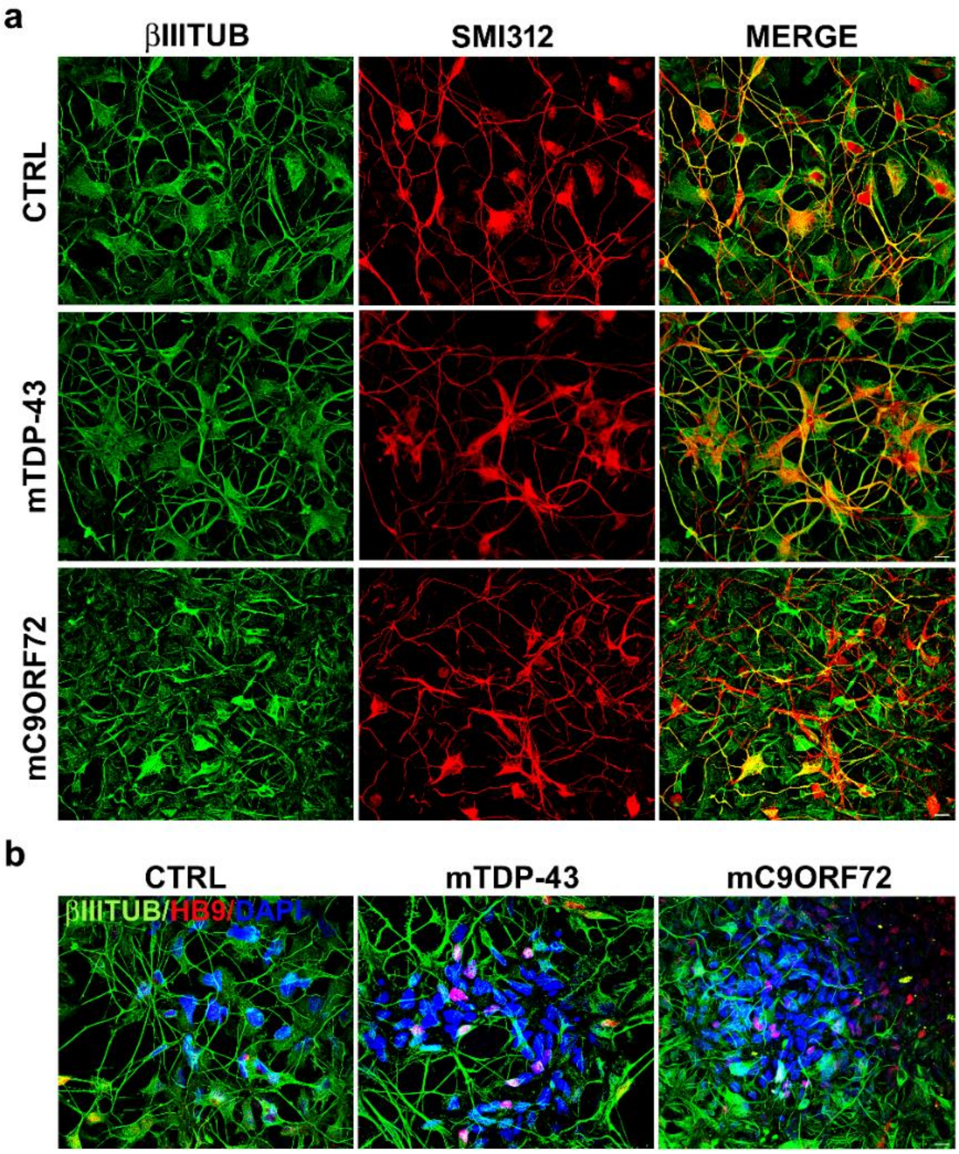
